# Genetic Tools for Conservation of Keystone Neotropical Raptors

**DOI:** 10.1101/2025.07.25.664441

**Authors:** Diego N. De Panis, Julián Padró

## Abstract

Neotropical raptors are among the most threatened avian groups, facing increasing extinction risks due to habitat loss and human persecution. Despite their importance for ecosystem stability, basic data on their distribution, abundance, and genetic diversity remain scarce. To address these gaps, we assembled and annotated the complete mitochondrial genomes of eight high-priority raptor species from the Neotropics, including the endangered Chaco Eagle (*Buteogallus coronatus*), Harpy Eagle (*Harpia harpyja*), and Rufous-tailed Hawk (*Buteo ventralis*). Mitogenome sizes ranged from 17,848 to 20,449 bp, with consistent gene content, and a Control Region architecture common in Falconidae and Accipitridae. Phylogenetic analyses provided strong support for most relationships, highlighting the value of mitogenomic data for phylogeographic studies. We further designed *in silico* metabarcoding primers for environmental DNA applications. Primers targeting the 12S rRNA gene and a mini-barcode for the Harpy Eagle’s Control Region showed high resolution using short, conserved sequences ideal for combining degraded DNA with next-generation sequencing. These resources enable evolutionary research and non-invasive biodiversity monitoring in difficult-to-survey habitats. Overall, our study provides essential genetic tools for monitoring and protecting these ecologically vital yet threatened birds across the Americas.

## Introduction

As apex predators, Neotropical raptors play a critical ecological role across the diverse landscapes of southern North America, Central America and South America: from lowland rainforests to the Patagonian steppes and the high Andean mountains (Sarasola et al., 2018). Comprising nearly one-third of all diurnal raptors (97 species comprising three orders: Accipitriformes, Falconiformes, and Cathartiformes), the Neotropical region harbors iconic species such as the Harpy Eagle (*Harpia harpyja*), the Crested Eagle (*Morphnus guianensis*) and the Chaco Eagle (*Buteogallus coronatus*). These keystone species are essential for regulating prey populations and maintaining ecosystem stability. However, their specialized traits, such as expansive home ranges, large body sizes and low reproductive rates, make them especially vulnerable to human pressures (Bildstein, 2017; Buechley et al., 2019). Nearly 25% of Neotropical raptors are now classified as being of conservation concern by The International Union for Conservation of Nature (IUCN): three Critically Endangered, three Endangered, seven Vulnerable and ten Near Threatened (Birdlife International 2025). Primary threats to their persistence include direct persecution, environmental contaminants, and especially habitat destruction (Sarasola et al. 2018). Despite this, basic knowledge gaps persist regarding their molecular diversity, distribution, abundance, and population trends for most species, particularly those inhabiting remote or inaccessible regions of the Neotropics (McClure et al., 2018; Buechley et al., 2019). Researchers have recently called for increased monitoring efforts and standardized surveys of wild populations (e.g., Perrig et al., 2019; Gousy-Leblanc et al., 2021; McClure et al., 2023, 2025). However, traditional survey methods are often inadequate for detecting elusive predators, which not only occur at low densities but also inhabit challenging environments such as dense forests, highlighting the need for innovative monitoring techniques.

A novel approach is to harness DNA from the environment (eDNA) to study wild populations without the need for direct observation. By coupling eDNA with metabarcoding libraries and next-generation sequencing, researchers can detect genetic material shed by organisms through skin cells, pellets, saliva, feces, or other biological traces left in water, soil, or air (Deiner et al., 2017; Ruppert et al., 2019; Clare et al., 2022). By analyzing informative markers, researchers can fill critical gaps in monitoring programs, while eliminating the need for invasive sampling, labor-intensive fieldwork, or reliance on expert opinion (e.g., Bourbour et al., 2024). This approach can provide data on species distribution, diversity, gene flow, and relative abundance, enabling more precise and cost-effective monitoring programs (Nakajima and Tsuri, 2024; Ai et al., 2025). Moreover, recent studies have shown that combining eDNA techniques with traditional methods can significantly enhance avian ecological research (Padró, 2024). However, the effectiveness of eDNA metabarcoding for biodiversity monitoring depends on the availability of comprehensive genetic reference databases for target species. Commonly used genetic markers, such as mitochondrial cytochrome *c* oxidase subunit I (COI), cytochrome *b* (CytB) and 12S rRNA, are critical for species identification but require well-curated sequences (Deiner et al., 2017). In addition, mitochondrial genes can provide insights into phylogenetic relationships, adaptations and cryptic diversity (De Panis et al., 2021). However, genetic resources for many raptor species remain limited, especially for those at high risk of extinction (Perrig et al., 2019; Gousy-Leblanc et al., 2021).

To address these gaps, we characterized the complete mitochondrial genomes of three threatened and five near-threatened Neotropical raptors. These include the endangered Chaco Eagle (*Buteogallus coronatus*), one of the rarest and most threatened eagles in South America. It inhabits open and semi-open lowland environments across southern Brazil, Bolivia, Paraguay, and central Argentina (Do, 2020). With an estimated global population of fewer than two thousand mature individuals, the species is severely affected by habitat loss due to deforestation, hunting, and electrocution on power lines (BirdLife International, 2024; Sarasola et al., 2020). A critical research priority is to determine whether genetic differentiation exists across its distribution, as such information is essential for guiding effective conservation strategies (Canal et al., 2017). The Rufous-tailed Hawk (*Buteo ventralis*) is also considered one of the rarest raptors in the Americas (Rivas-Fuenzalida et al., 2024). Currently, the species is largely confined to the temperate forests and adjacent open woodlands of southern Chile and Argentina. This medium-sized *Buteo* faces increasing threats from large-scale logging, the conversion of native forests into pine plantations, and persecution by ranchers. Classified as Vulnerable, the global population is estimated at fewer than one thousand mature individuals (BirdLife International, 2016). Yet, population sizes and habitat use remain poorly understood, highlighting the urgent need to assess its presence and abundance across the species’ distribution range (Rivas-Fuenzalida et al., 2024). The Harpy Eagle (*Harpia harpyja*), widely regarded as the world’s most powerful raptor, ranges from southern Mexico through Central America to the Amazon Basin and the Atlantic Forest, where it dominates the upper canopy of tropical rainforests. However, its distribution is patchy, with limited presence in the Atlantic Forest, as well as in Mexico, Guatemala, Belize, Costa Rica, and Ecuador (Schulenberg, 2020; BirdLife International, 2021). Wild populations are currently in decline, driven by large-scale deforestation, habitat fragmentation, and direct persecution. With an estimated global population of 100,000-250,000 mature individuals, the species is considered Vulnerable (BirdLife International 2021). However, given its low detectability its true status in many areas remains poorly understood (Schulenberg 2020). Moreover, the Harpy Eagle exhibits remarkably high mitochondrial genetic diversity, with most haplotypes unique to specific regions, and geographic differentiation both within South America and between Central and South America, highlighting the need for additional surveys to inform reintroduction efforts and conservation planning (Lerner et al., 2009).

To advance the conservation of threatened Neotropical raptors, this study leverages whole-genome sequencing to assemble and annotate complete mitochondrial genomes of eight high-priority species. We developed eDNA metabarcoding markers to help overcome critical limitations in species monitoring, clarify taxonomic uncertainties, and enhance phylogeographic and evolutionary analyses. This work lays the groundwork for integrating advanced molecular approaches into long-term strategies to protect these ecologically vital yet increasingly imperiled birds of prey.

## Material and Methods

### Assembly and Annotation

We assembled and annotated the complete mitochondrial genomes of the following keystone Neotropical raptors using whole-genome sequencing (WGS) data: *Buetogallus (*formerly *Harpyhaliaetus) coronatus* (SRR17853877), *Harpia harpyja* (SRR25728242/6/7), *Buteo ventralis* (SRR17454564), *Falco deiroleucus* (SRR24451039), *Buteogallus solitarius* (SRR17835758), *Cryptoleucopteryx plumbea* (SRR19616483), and *Spizaetus ornatus* (SRR18186383). In addition, we annotated the mitogenome of *Morphnus guianensis* (bMorGui1.MT.20240212) assembled from the whole-genome sequence data (GCA_045345515.1). For Illumina data, we used the assemblers NOVOPlasty v4.3.3 (Dierckxsens et al., 2017) and GetOrganelle v1.7.5 (Jin et al., 2020), selected for their efficiency in recovering circular mitogenomes from short-read sequences. PacBio HiFi long reads were assembled with MitoHiFi v3.2.2 (Uliano-Silva et al., 2023), leveraging HiFi accuracy to resolve highly contiguous genomes. Assembler algorithms were seeded with the longest available mitochondrial sequence per species, prioritizing the control region (CR) when present (Table S1). The resulting assemblies were annotated with MITOS v2 (Bernt et al., 2013) under the vertebrate mitochondrial genetic code, using RefSeq 89 Metazoa as a reference. Annotations were visualized as organelle genome maps with OGDRAW v1.3.1 (Greiner et al., 2019). To confirm species identity and exclude potential contamination, we conducted *in silico* validation using BLASTn (Chen et al., 2015) on informative markers (e.g., COI, CR, CytB) against NCBI records and verified sequence identity against published references for each species. All protein-coding genes (PCGs), rRNAs, tRNAs, and the CR were manually curated.

### Phylogenetic Analysis

To test the phylogenetic signal of our mitogenome dataset, we conducted comprehensive analyses incorporating 29 additional mitogenome sequences of diurnal raptor species (representing each genus) available from GenBank (Table S2). Our taxonomic sampling included a total of 37 raptor species representing Falconidae (5), Cathartidae (5), Sagittariidae (1), Pandionidae (1), and Accipitridae (25), plus nine outgroup taxa from Galliformes and Anseriformes. All analyses were performed using PhyloSuite v1.2.3 bioinformatics pipeline (Zhang et al., 2020). We standardized mitogenome annotations and performed sequence alignments with MAFFT v7.526 (Rozewicki et al., 2019) employing the codon-aware mode for PCGs and standard mode for RNAs. PCG alignments were subsequently refined with MACSE v2.04 (Ranwez et al., 2011), and ambiguously aligned regions were filtered with GBLOCKS v0.91b (Talavera and Castresana 2007) using conservative parameters (minimum block length = 10; gaps allowed). ModelFinder (Kalyaanamoorthy et al., 2017) was implemented to determine optimal partition schemes and substitution models through Bayesian Information Criterion scores, with unlinked branch lengths across partitions. We used the informative mitochondrial gene set (13 PCGs + 2 rRNAs) for the final phylogenetic reconstruction (De Panis et al., 2021), employing the maximum likelihood approach of IQ-TREE v2.4.0 (Nguyen et al., 2015) with 10,000 Ultra Fast Bootstraps with 1,000 iterations and 1,000 replicates under a partitioned model.

### Metabarcode Assessment

To develop taxon-specific metabarcoding primers for Neotropical raptor eDNA monitoring, we implemented a bioinformatics pipeline for *in silico* primer design (Ficetola et al., 2010). First, we compiled a reference database using BLASTn (Chen et al., 2015) to query our annotated COI, CytB, and 12S sequences (loci with established discriminatory power). We retained all Neotropical sympatric raptor species (one representative per species) from GenBank to compile the target database, reaching a total of 134 sequences from 57 species (Table S2). These curated sequences were analyzed in the EcoPrimers package (Riaz et al., 2011) to develop potential barcode primers with taxonomic resolution for eDNA metagenomic applications. We used standard parameters: a maximum of three mismatches allowed, minimized amplicon lengths (prioritizing <300 bp for degraded eDNA and NGS compatibility), and optimization for GC content and melting temperature uniformity. Candidate primers were further filtered to minimize dimer formation and maximize binding coverage (Bc), defined as the proportion of target taxa with perfect primer matches. Additionally, we used the EcoPCR package (Ficetola et al., 2010) to further validate *in silico* specificity against the reference database (Bellemain et al., 2010). We also applied this pipeline to all 22 known Harpy Eagle CR haplotypes (∼400 bp; Table S2) to develop eDNA minibarcoding primers for intraspecific resolution (<200 bp), enabling potential haplotype-level population monitoring (Lerner et al., 2009). This approach prioritized primer sets that balance amplification efficiency with taxonomic discrimination, ensuring suitability for eDNA applications where template DNA is often fragmented and present in low quantities.

## Results and Discussion

Neotropical raptors face significant conservation challenges, with most species either threatened or lacking adequate population data. In this study, we present the complete mitochondrial genome assemblies and annotations for eight emblematic raptor species of the Neotropics, including three threatened species: the Chaco Eagle (*Buteogallus coronatus*), Harpy Eagle (*Harpia harpyja*), and Rufous-tailed Hawk (*Buteo ventralis*). We also included five near-threatened and data-deficient taxa: the Orange-breasted Falcon (*Falco deiroleucus*), Solitary Eagle (*Buteogallus solitarius*), Plumbeous Hawk (*Cryptoleucopteryx plumbea*), Ornate Hawk-Eagle (*Spizaetus ornatus*) and the Crested Eagle (*Morphnus guianensis*). In addition, we assessed the phylogenetic resolution of mitogenome sequences and developed *in silico* metabarcoding primers for environmental DNA applications, providing crucial resources for conservation genetics and biodiversity monitoring.

The mitochondrial genome size of the eight keystone Neotropical raptors ranged from 17,848 bp to 20,449 bp, with the Orange-breasted Falcon having the smallest genome and the Ornate Hawk-Eagle the largest one. All genomes contained the full set of 37 genes typically found in vertebrates, including 22 transfer RNAs (tRNAs), 2 ribosomal RNAs (rRNAs), 13 protein-coding genes (PCGs), and a Control region (CR; Fig. 1). These genetic resources not only expand the mitogenomic representation of eight raptor species, but also include four genera previously unrepresented in public databases (Fig. 2). Moreover, they provide species-identification markers that were previously unavailable, including COI for six of the eight species, 12S and CR for five, as well as CytB for one (Table S1). The overall nucleotide composition of all mitogenomes was comparable (A ≈ 31.2%, C ≈ 31.1%, G ≈ 13.7%, T ≈ 24%), exhibiting a slight A+T bias (Fig. S1), in line with the general tendency of most mitochondrial genomes of raptors (De Panis et al., 2021). The Control region ranged from 1,046 bp (Orange-breasted Falcon) to 2,777 bp (Chaco Eagle). Unlike the typical ancestral gene order seen in species such as *Gallus gallus*, or New World vultures (De Panis et al., 2021) where the Control region is located between tRNA-Glu and tRNA-Phe, the CR of the newly characterized raptor species is positioned between tRNA-Thr and tRNA-Pro (Fig. 1). In addition, the species exhibited the presence of a variable pseudo Control region between tRNA-Glu and tRNA-Phe, where the ancestral arrangement includes the Control region (Fig. 1). This is consistent with the mitochondrial architecture observed in many Falconidae and Accipitridae species such as *Falco peregrinus* (Mindell et al., 1998) and *Buteo buteo* (Haring et al., 2001). However, while our assembly and annotation pipeline successfully recovered complete mitochondrial genomes with all Control regions matching expected references, some technical limitations may remain. For instance, short-read Illumina sequencing may introduce errors in highly repetitive regions, potentially affecting length estimates and tandem repeat detection. Thus, experimental validation (e.g., long-read sequencing or PCR confirmation) of these regions would further strengthen these results.

**Figure 1.**
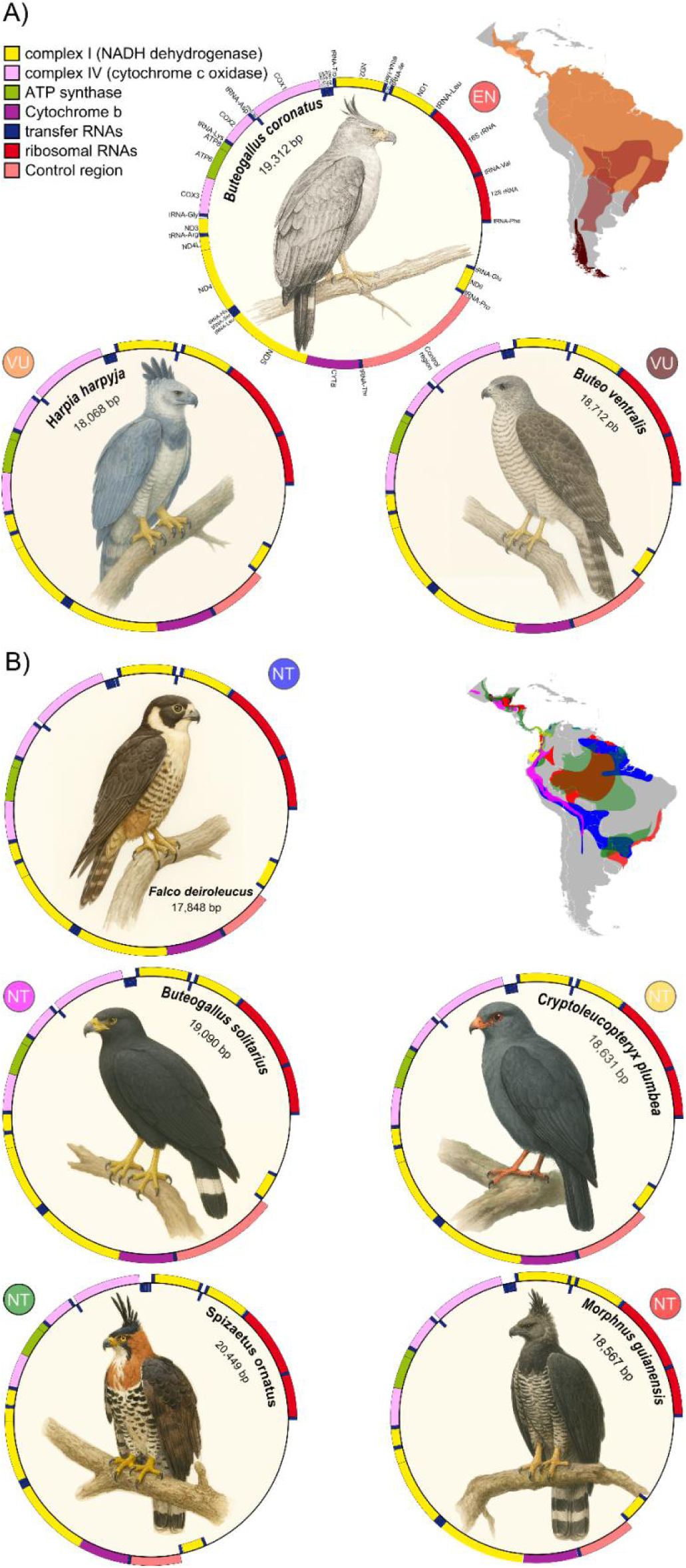
Physical map of the complete mitochondrial genomes of three threatened (A) and five near-threatened (B) top avian predators of the Neotropics. The conservation status of each species is indicated by colored circles and their distributional range are shown on the map (color-coded).

**Figure 2.**
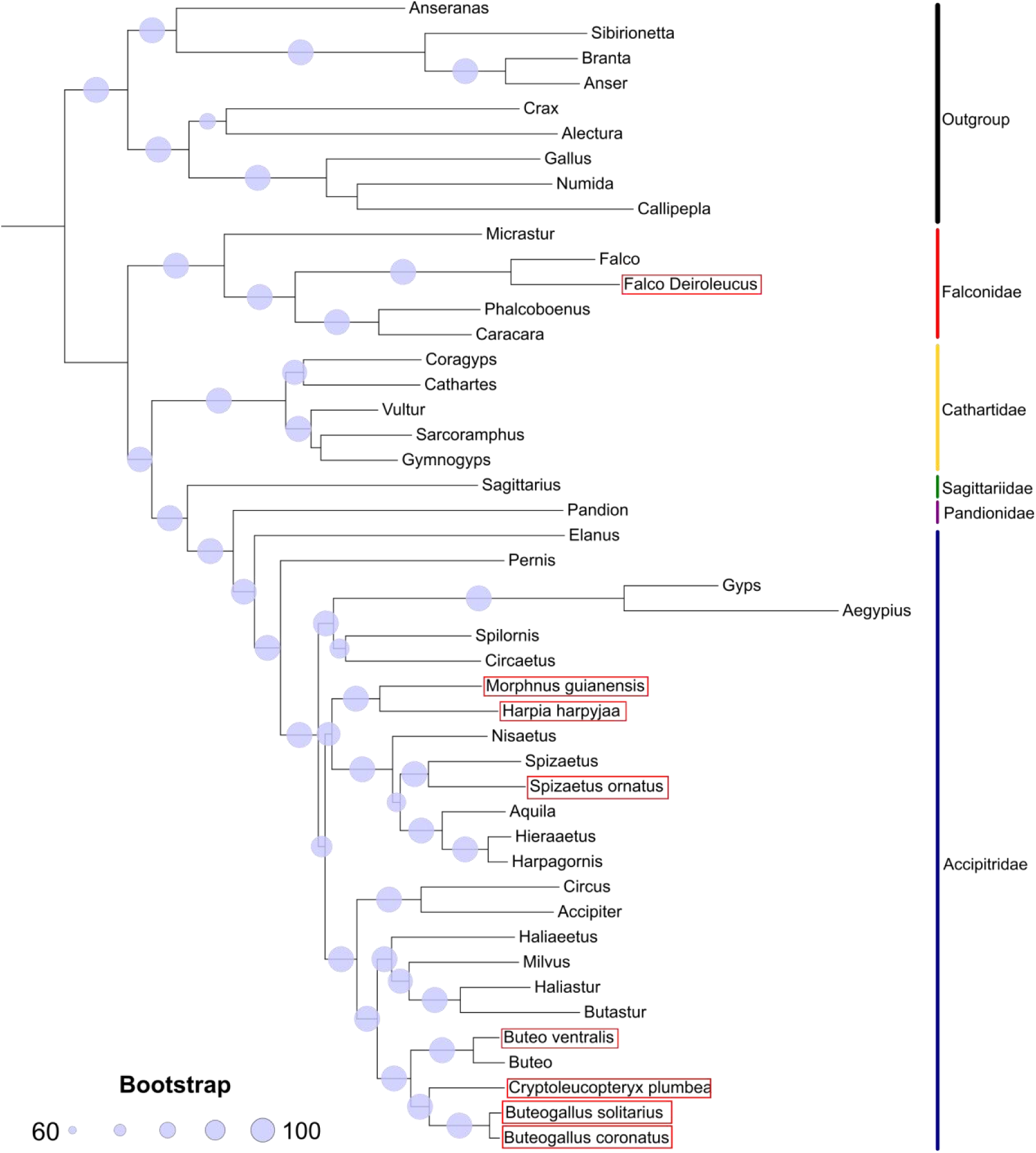
Phylogenetic relationships of the Orange-breasted falcon (*Falco deiroleucus*) Crested Eagle (*Morphnus guianensis*), Harpy Eagle (*Harpia harpyja*), Ornate Hawk-Eagle (*Spizaetus ornatus*), Rufous-tailed Hawk (*Buteo ventralis*), Plumbeous Hawk (*Cryptoleucopterix plumbea*), Solitary Eagle (*Buteogallus solitarius*) and Chaco Eagle (*Buteogallus coronatus*), based on a maximum likelihood tree of mitogenomic sequences.

To test the phylogenetic resolution of the assembled mitogenomes, we analyzed the systematics of taxonomically challenging groups such as Accipitridae, which encompasses several cosmopolitan species of hawks, Old World vultures and eagles. Overall, our phylogenetic reconstruction of raptor species was supported by most of the nodes with bootstrap values >95. An exception to this pattern was observed in the Cathartidae family, which, although not a focal group in our study, was included in the analysis and exhibited limited phylogenetic resolution, with a bootstrap value of 50 for the sister relationship between *Sarcoramphus* and *Gymnogyps* (Fig. 2). This group has previously been described as challenging to resolve using a combination of nuclear and mitochondrial loci (Johnson et al., 2016). Nonetheless, recent phylogenomic analyses using ultraconserved elements (UCEs) recovered *Sarcoramphus* and *Vultur* as sister taxa with strong support (Catanach et al., 2025). Within falcons, our analysis confirmed the traditional classification placing caracaras (*Phalcoboenus* and *Caracara*) in a distinct clade from true falcons (Fuchs et al., 2015). Both *Morphnus* and *Harpia* were placed as sister taxa, consistent with the traditional subfamily Harpiinae (*Harpia, Morphnus, Harpyopsis* and *Macheiramphus*), showing a similar branching order described in previous phylogenies (Lerner and Mindell, 2005; Mindell et al., 2018), which place the subfamily as sister to Aquilinae (eagle species, including our target *Spizaetus ornatus*). However, this branching pattern differs from recent UCE-based reconstruction showing Harpiinae as a sister group to Buteoninae + Accipitrinae (Catanach et al., 2025). Finally, our phylogenomic analysis recovered the subfamily Buteoninae composed of two major subclades corresponding to the tribes Milvini (*Haliaeetus, Milvus,* and *Haliastur*) and Buteonini (*Buteo, Cryptoleucopteryx, Buteogallus*). Yet, our analysis placed *Butastur* in Milvini instead of the traditionally Buteonini clade, likely due to incomplete taxon sampling, emphasizing the need for more comprehensive mitogenomic data across raptor species (do Amaral et al., 2009; Mindell et al., 2018; Catanach et al., 2025).

Using our assembled mitogenomes along with reference sequences of raptor species from the tropical Americas, we developed *in silico* primers for metabarcoding applications in biodiversity monitoring. We first focused on the standard barcode for animal species, the Cytochrome *c* Oxidase subunit I (Ratnasingham and Hebert, 2007), and also evaluated the Cytochrome *b*, a traditional marker for raptor phylogenetics (e.g., Seibold and Helbig, 1995; Wink, 1995; Wink et al., 1998), which has proven effective for species identification in some taxa (Deiner et al., 2017; Ruppert et al., 2019). Our best candidate barcodes for both loci with a reasonable fragment size (<200 bp; Fig. 3) achieved a discrimination power of 96%, failing only to distinguish between *Buteo ventralis* and *Buteo jamaicensis.* However, as these species are allopatric, geographic separation minimizes the risk of misidentification. Nonetheless, both COI and CytB proved suboptimal for metabarcoding, due to high variability leading to either long fragments incompatible with degraded eDNA or short fragments lacking conserved primer-binding sites (Deagle et al., 2014). Primer design for these loci required numerous degenerate bases (Table 1), increasing the risk of non-specific amplification (additional results in Table S3).

**Figure 3.**
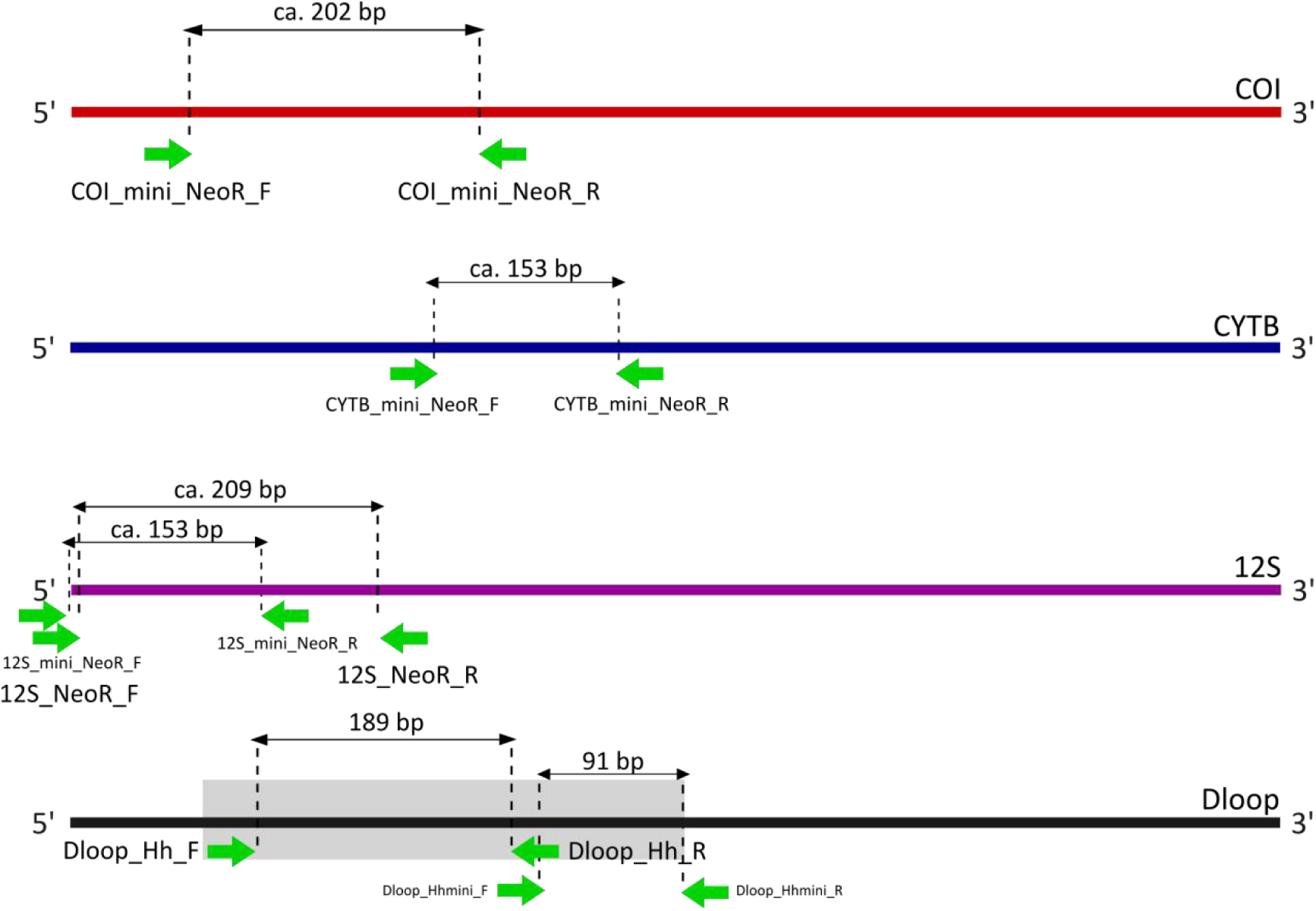
Relative positions of primer binding sites across the complete mitochondrial genes and the Control Region (the gray box indicates the characterized polymorphic region in *H. harpyja*). The average total length of each barcode region is shown.

**Table 1.**
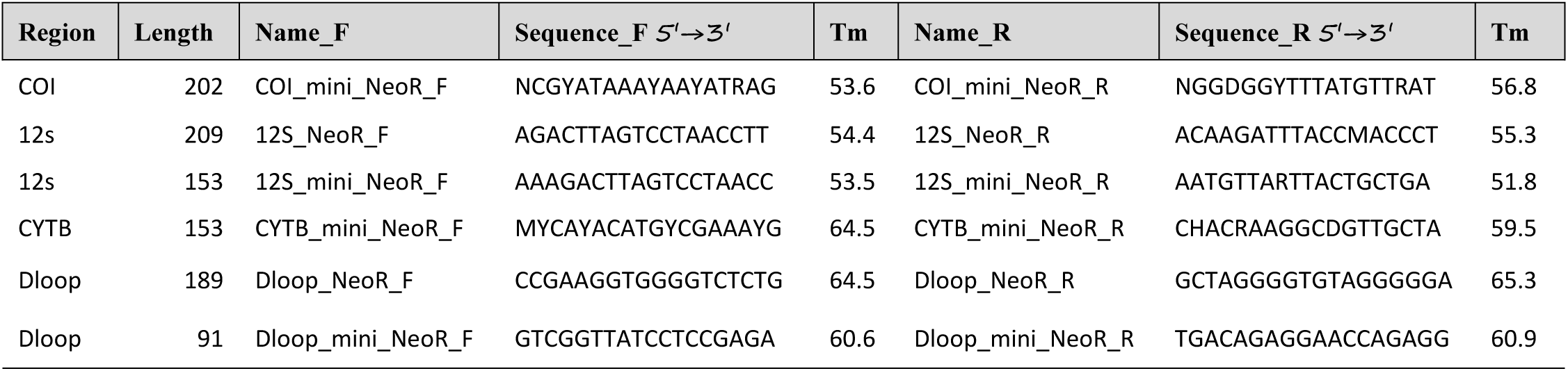
Best primer pairs for targeted barcode amplification across mitochondrial markers in keystone Neotropical raptor species.

Our most effective barcode candidate was the 12S ribosomal RNA gene, which is well-suited for eDNA studies using degraded samples. The longer (209 bp) version exhibited highly conserved primer-binding regions (Table 1; Fig. 3), identifying all target species of conservation concern. This marker achieved an overall 94% discrimination power, failing only to distinguish between *Buteo swainsoni* and *Buteo brachyurus* (largely allopatric during the breeding season), and between the allopatric pair *Pseudastur albicollis* and *Pseudastur polionotus* (BirdLife International, 2025). Our mini-barcode version spanning 153 bp required only one degenerate base in the reverse primer (Table 1) and showed similar resolution, with an additional ambiguity between *Buteogallus aequinoctialis* and *Buteogallus anthracinus* which are also largely allopatric. Designed for ultra-short targets, this mini-barcode is expected to enhance species detection in samples containing trace or highly degraded DNA, such as soil, decomposed remains, or even air samples (Epp et al., 2012; Clare et al., 2022; Goray et al., 2024). The choice of the 12S rRNA gene aligns with its growing status as a standard marker for animal metabarcoding, due to the unique features of its RNA secondary structure (Riaz et al., 2011; Clarke et al., 2014; Deagle et al., 2014). This gene alternates between conserved and variable regions, with stable stem structures providing consistent primer-binding sites flanked by hypervariable loops that facilitate species-level discrimination (Kocher et al., 2017).

Finally, we developed species-specific barcode primers targeting the Control Region of the Harpy Eagle. The longer fragment (189 bp) successfully distinguished 20 of the 22 known haplotypes (Lerner et al., 2009) but failed to differentiate between haplotypes 3 and 20, as the diagnostic polymorphism lies just beyond the 3′ end of the amplified region (at the extreme end of Domain I). Despite this limitation, the two haplotypes seem to occur in distinct geographic regions: haplotype 3 was found in Peru, while haplotype 20 was found in northern South America and Central America (Lerner et al., 2009). To overcome this limitation and enable identification of all 22 haplotypes using shorter fragments compatible with eDNA metabarcoding approaches, we designed a mini-barcode primer set (91 bp). These primers target highly conserved regions flanking the key diagnostic polymorphism, allowing for full resolution of all haplotypes sourced from degraded DNA samples (Table 1; Fig. 3).

In conclusion, the newly assembled mitochondrial genomes of these keystone Neotropical raptor species provide a valuable resource for advancing avian conservation genetics. These data help resolve phylogenetic relationships in taxonomically complex groups and support the molecular identification of species, lineages, and their geographic origins. By enabling detection of spatial haplotype patterns, they inform inferences about gene flow and the delineation of potential biological corridors, which are essential for conservation planning. Mitogenomes also offer insight into local adaptation and divergence, contributing to the identification of evolutionarily significant units. Finally, the development of metabarcoding primers expands the utility of these data for non-invasive biodiversity monitoring via environmental DNA, particularly in remote or logistically challenging ecosystems.

Together, these molecular tools enhance our capacity to study and protect some of the Neotropics’ most iconic raptors amid accelerating environmental change.

## Acknowledgments

This study was supported by the National Scientific and Technical Research Council of Argentina (CONICET). We thank Juan Manuel Grande for his review of an early draft of the manuscript. The mitochondrial genome data for *Morphnus guianensis* was obtained through GenomeArk (Vertebrate Genomes Project) after prior consultation. Details of data analyses and accession numbers for sequences retrieved from GenBank have been uploaded as part of the Supplementary Material. Additionally, mitogenomic sequences generated in this study will be deposited in GenBank, the NIH genetic sequence database.

## Notes

### Competing Interest Statement

The authors have declared no competing interest.

