## Supplementary material for "Genetic Tools for Conservation of Keystone Neotropical Raptors"

Table S1. GenBank accession numbers of mitochondrial sequences used for mitogenome assembly (seed) and metabarcoding analyses.

| Species / Marker | COI | 12S | CYTB | CR / SEED |
|---|---|---|---|---|
| Accipiter bicolor | FJ027014.1 | - | - | |
| Busarellus nigricollis | FJ027235.1 | GQ264649.1 | GQ264810.1 | |
| Buteo albigula | - | GQ264601.1 | GQ264774.1 | |
| Buteo albonotatus | JN801523.1 | GQ264602.1 | GQ264775.1 | |
| Buteo brachyurus | MT456669.1 | GQ264608.1 | GQ264781.1 | |
| Buteo jamaicensis | KR017962.1 | GQ264618.1 | GQ264784.1 | |
| Buteo lineatus | KR017961.1 | GQ264623.1 | GQ264788.1 | |
| Buteo nitidus | DQ433339.1 | GQ264629.1 | GQ264794.1 | |
| Buteo platypterus | DQ432787.1 | GQ264632.1 | GQ264796.1 | |
| Buteo regalis | KF525369.2 | GQ264637.1 | GQ264802.1 | |
| Buteo solitarius | - | GQ264643.1 | GQ264804.1 | |
| Buteo swainsoni | DQ433391.1 | GQ264645.1 | EU583345.1 | |
| Buteo ventralis | - | - | - | AY213024 |
| Buteogallus aequinoctialis | JQ174209.1 | GQ264597.1 | GQ264770.1 | |
| Buteogallus anthracinus | DQ432784.1 | GQ264606.1 | GQ264779.1 | |
| Buteogallus meridionalis | FJ027237.1 | GQ264628.1 | GQ264792.1 | |
| Buteogallus schistaceus | - | GQ264685.1 | EU583373.1 | |
| Buteogallus solitarius | - | GQ264659.1 | GQ264820.1 | GQ264659.1 |
| Buteogallus subtilis | - | GQ264644.1 | EU583331.1 | |
| Buteogallus urubitinga | FJ027240.1 | DQ148351.1 | GQ264809.1 | |
| Caracara cheriway | - | NC_044673.1 | - | |
| Caracara plancus | - | NC_044672.1 | - | |
| Cryptoleucopterix plumbea | - | GQ264680.1 | GQ264841.1 | GQ264680.1 |
| Falco columbarius | AY666522.1 | NC_025579.1 | NC_025579.1 | |
| falco deiroleucus | - | - | EU233056.1 | KM876051.1 |
| Falco femoralis | FJ027579.1 | - | U83310.1 | |
| Falco mexicanus | AY666553.2 | - | EU233076.1 | |
| Falco peregrinus | AB842766.1 | NC_000878.1 | JQ282801.1 | |
| Falco rufigularis | JQ174836.1 | - | - | |
| Falco sparverius | KR017978.1 | NC_008547.1 | - | |
| Geranoaetus albicaudatus | DQ433389.1 | GQ264599.1 | GQ264773.1 | |
| Geranoaetus melanoleucus | FJ027626.1 | GQ264653.1 | GQ264813.1 | |
| Geranoaetus poecilochrous | - | - | EU583340.1 | |
| Geranoaetus polyosoma | FJ027251.1 | GQ264634.1 | GQ264798.1 | |
| Geranospiza caerulescens | KM894332.1 | GQ264651.1 | GQ264811.1 | |
| harpia harpyja | - | - | AJ604495.1 | GQ917189-211.1 |
| Harpyhaliaetus coronatus | JN801708.1 | GQ264654.1 | GQ264815.1 | |
| Ictinia mississippiensis | FJ027681.1 | GQ264660.1 | GQ264821.1 | |

| | | | | |
|---|---|---|---|---|
| Ictinia plumbea | FJ027683.1 | GQ264661.1 | GQ264822.1 | |
| Leucopternis kuhli | JN801774.1 | GQ264669.1 | EU583358.1 | |
| Leucopternis lacernulatus | JN801775.1 | DQ148340.1 | GQ264832.1 | |
| Leucopternis melanops | JQ175259.1 | GQ264673.1 | GQ264836.1 | |
| Leucopternis semiplumbeus | JQ175261.1 | GQ264686.1 | GQ264846.1 | |
| Morphnarchus princeps | JQ175260.1 | GQ264683.1 | GQ264843.1 | |
| Morphnus guianensis | - | - | AJ604496.1 | GQ917209.1 |
| Pandion haliaetus | NC_008550.1 | - | - | |
| Parabuteo leucorrhous | FJ027243.1 | GQ264688.1 | GQ264849.1 | |
| Parabuteo unicinctus | AY666176.1 | GQ264693.1 | GQ264853.1 | |
| Phalcoboenus australis | - | NC_031897.1 | - | |
| Pseudastur albicollis albicollis | JN801773.1 | GQ264662.1 | GQ264826.1 | |
| Pseudastur occidentalis | - | GQ264676.1 | EU583364.1 | |
| Pseudastur polionotus | - | GQ264682.1 | GQ264842.1 | |
| Rostrhamus sociabilis | FJ028214.1 | GQ264695.1 | GQ264854.1 | |
| Rupornis magnirostris | FJ027245.1 | GQ264625.1 | GQ264790.1 | |
| Spizaetus isidori | - | - | AJ812238.1 | |
| Spizaetus ornatus | KU842344.1 | - | AJ604508.2 | EF459593.1 |
| Spizaetus tyrannus | KU842345.1 | NC_052803.1 | AJ604510.2 | |

Table S2. GenBank accession numbers for the complete metogenomes of bird species used in phylogenetic analyses.

| Family | Species name | Accession |
|---|---|---|
| Anseranatidae | Anseranas semipalmata | NC_005933.1 |
| Anatidae | Sibirionetta formosa | NC_015482.1 |
| Anatidae | Branta canadensis | NC_007011.1 |
| Anatidae | Anser cygnoides | NC_023832.1 |
| Cracidae | Crax rubra | NC_024618.1 |
| Megapodiidae | Alectura lathami | NC_007227.1 |
| Phasianidae | Gallus gallus spadiceus | NC_040902.1 |
| Numididae | Numida meleagris | NC_034374.1 |
| Odontophoridae | Callipepla squamata | NC_029340.1 |
| Falconidae | Micrastur gilvicollis | NC_008548.1 |
| Falconidae | Falco peregrinus | NC_000878.1 |
| Falconidae | Phalcoboenus australis | NC_031897.1 |
| Falconidae | Caracara plancus | NC_044672.1 |
| Cathartidae | Coragyps atratus | MN720440.1 |
| Cathartidae | Cathartes aura | NC_007628.1 |
| Cathartidae | Vultur gryphus | MZ223429 |
| Cathartidae | Sarcoramphus papa | MN720442.1 |
| Cathartidae | Gymnogyps californianus | BK059163 |
| Sagittariidae | Sagittarius serpentarius | NC_023788.1 |
| Pandionidae | Pandion haliaetus | NC_008550.1 |
| Accipitridae | Elanus caeruleus | OK662584.1 |
| Accipitridae | Pernis ptilorhynchus | LC541458.1 |
| Accipitridae | Gyps fulvus | NC_036050.1 |
| Accipitridae | Aegypius monachus | KF682364.1 |
| Accipitridae | Spilornis cheela | NC_015887.1 |
| Accipitridae | Circaetus pectoralis | NC_052805.1 |
| Accipitridae | Nisaetus nipalensis | NC_007598.1 |
| Accipitridae | Spizaetus tyrannus | NC_052803.1 |
| Accipitridae | Aquila heliaca | NC_035806.1 |
| Accipitridae | Hieraaetus pennatus | MK294165.1 |
| Accipitridae | Harpagornis moorei | MK294166.1 |
| Accipitridae | Circus melanoleucos | NC_035801.1 |
| Accipitridae | Accipiter gentilis | NC_011818.1 |
| Accipitridae | Haliaeetus albicilla | NC_040858.1 |
| Accipitridae | Milvus migrans | NC_038195.1 |
| Accipitridae | Haliastur indus | OP133375.1 |
| Accipitridae | Butastur indicus | NC_032362.1 |
| Accipitridae | Buteo buteo | NC_003128.3 |

Table S3. Additional primers developed *in silico* for three mitochondrial markers across keystone neotropical raptors.

| **Marker** | **COI** | | | | | | | | |
|---|---|---|---|---|---|---|---|---|---|
| **Primer 1** | **Primer 2** | **tm1** | **tm2** | **Quality** | ***Bc* index** | ***Bs* index** | **Min length** | **Max length** | **Av. length** |
| ACGCATAAACAACATAAG | TGGGGGTTTTATGTTGAT | 48.9 | 51.5 | GG | 0.98 | 0.96 | 202 | 202 | 202 |
| CATAAACAACATAAGCTT | TGGGGGTTTTATGTTGAT | 45.9 | 51.5 | GG | 0.98 | 0.96 | 199 | 199 | 199 |
| ATAAACAACATAAGCTTC | TGGGGGTTTTATGTTGAT | 45.6 | 51.5 | GG | 0.98 | 0.96 | 198 | 198 | 198 |
| TAAACAACATAAGCTTCT | TGGGGGTTTTATGTTGAT | 46.6 | 51.5 | GG | 0.98 | 0.96 | 197 | 197 | 197 |
| AAACAACATAAGCTTCTG | TGGGGGTTTTATGTTGAT | 48.6 | 51.5 | GG | 0.98 | 0.96 | 196 | 196 | 196 |
| CATAAGCTTCTGACTACT | TGGGGGTTTTATGTTGAT | 48.7 | 51.5 | GG | 0.96 | 0.95 | 190 | 190 | 190 |
| AACAACATAAGCTTCTGA | TGGGGGTTTTATGTTGAT | 49.3 | 51.5 | GG | 0.93 | 0.95 | 195 | 195 | 195 |
| CAACATAAGCTTCTGACT | TGGGGGTTTTATGTTGAT | 50 | 51.5 | GG | 0.93 | 0.95 | 193 | 193 | 193 |
| GCATAAACAACATAAGCT | TGGGGGTTTTATGTTGAT | 48.8 | 51.5 | GG | 0.91 | 0.95 | 200 | 200 | 200 |
| AGCCTCCTCAACAGTAGA | TGGGGGTTTTATGTTGAT | 54.7 | 51.5 | GG | 0.96 | 0.91 | 148 | 148 | 148 |
| CGCATAAACAACATAAGC | TGGGGGTTTTATGTTGAT | 50.6 | 51.5 | GG | 0.91 | 0.83 | 201 | 201 | 201 |
| CTAGCCTCCTCAACAGTA | TGGGGGTTTTATGTTGAT | 53 | 51.5 | GG | 0.96 | 0.65 | 150 | 150 | 150 |
| TAGCCTCCTCAACAGTAG | TGGGGGTTTTATGTTGAT | 53 | 51.5 | GG | 0.96 | 0.65 | 149 | 149 | 149 |
| CTCCTCAACAGTAGAAGC | TGGGGGTTTTATGTTGAT | 52.7 | 51.5 | GG | 0.96 | 0.61 | 145 | 145 | 145 |
| CTCAACAGTAGAAGCAGG | TGGGGGTTTTATGTTGAT | 53 | 51.5 | GG | 0.96 | 0.61 | 142 | 142 | 142 |
| **Marker** | **CYTB** | | | | | | | | |
| AGGCCAAATATCCTTCTG | TGCTGGGGTGAAGTTTTC | 52 | 55.9 | GG | 0.98 | 0.96 | 337 | 337 | 337 |
| CCAAATATCCTTCTGAGG | TGCTGGGGTGAAGTTTTC | 49.8 | 55.9 | GG | 0.98 | 0.96 | 334 | 334 | 334 |
| GCCAAATATCCTTCTGAG | TGCTGGGGTGAAGTTTTC | 50.6 | 55.9 | GG | 0.96 | 0.96 | 335 | 335 | 335 |
| AGGCCAAATATCCTTCTG | TGGGGTGAAGTTTTCTGG | 52 | 55.1 | GG | 0.94 | 0.96 | 334 | 334 | 334 |
| AGGCCAAATATCCTTCTG | CTGGGGTGAAGTTTTCTG | 52 | 53.8 | GG | 0.94 | 0.96 | 335 | 335 | 335 |
| CCAAATATCCTTCTGAGG | TGGGGTGAAGTTTTCTGG | 49.8 | 55.1 | GG | 0.94 | 0.96 | 331 | 331 | 331 |
| CCCACACATGCCGAAACG | CTACGAAGGCAGTTGCTA | 60.5 | 54.9 | GG | 0.98 | 0.92 | 153 | 153 | 153 |
| CCCACACATGCCGAAACG | GAAGGATATTTGGCCTCA | 60.5 | 52 | GG | 0.98 | 0.92 | 185 | 185 | 185 |
| CCACACATGCCGAAACGT | TACGAAGGCAGTTGCTAT | 59.5 | 53.9 | GG | 0.98 | 0.92 | 151 | 151 | 151 |
| CCACACATGCCGAAACGT | CTACGAAGGCAGTTGCTA | 59.5 | 54.9 | GG | 0.98 | 0.92 | 152 | 152 | 152 |
| CCACACATGCCGAAACGT | CCTACGAAGGCAGTTGCT | 59.5 | 57.8 | GG | 0.98 | 0.92 | 153 | 153 | 153 |
| CCACACATGCCGAAACGT | GAAGGATATTTGGCCTCA | 59.5 | 52 | GG | 0.98 | 0.92 | 184 | 184 | 184 |
| AGGCCAAATATCCTTCTG | GGTGAAGTTTTCTGGGTC | 52 | 53.8 | GG | 0.94 | 0.96 | 331 | 331 | 331 |
| CCAAATATCCTTCTGAGG | GGTGAAGTTTTCTGGGTC | 49.8 | 53.8 | GG | 0.94 | 0.96 | 328 | 328 | 328 |
| GCCAAATATCCTTCTGAG | GGTGAAGTTTTCTGGGTC | 50.6 | 53.8 | GG | 0.92 | 0.96 | 329 | 329 | 329 |
| ACATGCCAACGGAGCATC | TACGAAGGCAGTTGCTAT | 58.9 | 53.9 | GG | 0.94 | 0.92 | 106 | 106 | 106 |
| ACATGCCAACGGAGCATC | CTACGAAGGCAGTTGCTA | 58.9 | 54.9 | GG | 0.94 | 0.92 | 107 | 107 | 107 |
| ACATGCCAACGGAGCATC | CCTACGAAGGCAGTTGCT | 58.9 | 57.8 | GG | 0.94 | 0.92 | 108 | 108 | 108 |
| ACATGCCAACGGAGCATC | GAAGGATATTTGGCCTCA | 58.9 | 52 | GG | 0.94 | 0.92 | 139 | 139 | 139 |
| AGGCCAAATATCCTTCTG | TTGCTGGGGTGAAGTTTT | 52 | 55.1 | GG | 0.98 | 0.88 | 338 | 338 | 338 |

| | | | | | | | | | |
|---|---|---|---|---|---|---|---|---|---|
| AGGCCAAATATCCTTCTG | TTTGCTGGGGTGAAGTTT | 52 | 55.1 | GG | 0.98 | 0.88 | 339 | 339 | 339 |
| AGGCCAAATATCCTTCTG | GTTTGCTGGGGTGAAGTT | 52 | 56.2 | GG | 0.98 | 0.88 | 340 | 340 | 340 |
| CCAAATATCCTTCTGAGG | TTGCTGGGGTGAAGTTTT | 49.8 | 55.1 | GG | 0.98 | 0.88 | 335 | 335 | 335 |
| CCAAATATCCTTCTGAGG | TTTGCTGGGGTGAAGTTT | 49.8 | 55.1 | GG | 0.98 | 0.88 | 336 | 336 | 336 |
| CCAAATATCCTTCTGAGG | GTTTGCTGGGGTGAAGTT | 49.8 | 56.2 | GG | 0.98 | 0.88 | 337 | 337 | 337 |
| GGCCAAATATCCTTCTGA | TGCTGGGGTGAAGTTTTC | 52 | 55.9 | GG | 0.96 | 0.88 | 336 | 336 | 336 |
| GGCCAAATATCCTTCTGA | TTGCTGGGGTGAAGTTTT | 52 | 55.1 | GG | 0.96 | 0.88 | 337 | 337 | 337 |
| GGCCAAATATCCTTCTGA | TTTGCTGGGGTGAAGTTT | 52 | 55.1 | GG | 0.96 | 0.88 | 338 | 338 | 338 |
| GGCCAAATATCCTTCTGA | GTTTGCTGGGGTGAAGTT | 52 | 56.2 | GG | 0.96 | 0.88 | 339 | 339 | 339 |
| GCCAAATATCCTTCTGAG | TTGCTGGGGTGAAGTTTT | 50.6 | 55.1 | GG | 0.96 | 0.88 | 336 | 336 | 336 |
| GCCAAATATCCTTCTGAG | TTTGCTGGGGTGAAGTTT | 50.6 | 55.1 | GG | 0.96 | 0.88 | 337 | 337 | 337 |
| GCCAAATATCCTTCTGAG | GTTTGCTGGGGTGAAGTT | 50.6 | 56.2 | GG | 0.96 | 0.88 | 338 | 338 | 338 |
| CACATGCCGAAACGTACA | CTACGAAGGCAGTTGCTA | 56.6 | 54.9 | GG | 0.90 | 0.91 | 149 | 149 | 149 |
| CACATGCCGAAACGTACA | GAAGGATATTTGGCCTCA | 56.6 | 52 | GG | 0.90 | 0.91 | 181 | 181 | 181 |
| CTACGAAGGCAGTTGCTA | TACATGCCAACGGAGCAT | 54.9 | 57.3 | GG | 0.90 | 0.91 | 108 | 108 | 108 |
| GAAGGATATTTGGCCTCA | TACATGCCAACGGAGCAT | 52 | 57.3 | GG | 0.90 | 0.91 | 140 | 140 | 140 |
| AGGCCAAATATCCTTCTG | GCTGGGGTGAAGTTTTCT | 52 | 55.5 | GG | 0.94 | 0.87 | 336 | 336 | 336 |
| CCAAATATCCTTCTGAGG | CTGGGGTGAAGTTTTCTG | 49.8 | 53.8 | GG | 0.94 | 0.87 | 332 | 332 | 332 |
| CCAAATATCCTTCTGAGG | GCTGGGGTGAAGTTTTCT | 49.8 | 55.5 | GG | 0.94 | 0.87 | 333 | 333 | 333 |
| GGCCAAATATCCTTCTGA | TGGGGTGAAGTTTTCTGG | 52 | 55.1 | GG | 0.92 | 0.87 | 333 | 333 | 333 |
| CTGGGGTGAAGTTTTCTG | GGCCAAATATCCTTCTGA | 53.8 | 52 | GG | 0.92 | 0.87 | 334 | 334 | 334 |
| GCTGGGGTGAAGTTTTCT | GGCCAAATATCCTTCTGA | 55.5 | 52 | GG | 0.92 | 0.87 | 335 | 335 | 335 |
| GCCAAATATCCTTCTGAG | TGGGGTGAAGTTTTCTGG | 50.6 | 55.1 | GG | 0.92 | 0.87 | 332 | 332 | 332 |
| CTGGGGTGAAGTTTTCTG | GCCAAATATCCTTCTGAG | 53.8 | 50.6 | GG | 0.92 | 0.87 | 333 | 333 | 333 |
| GCCAAATATCCTTCTGAG | GCTGGGGTGAAGTTTTCT | 50.6 | 55.5 | GG | 0.92 | 0.87 | 334 | 334 | 334 |
| GGCCAAATATCCTTCTGA | GGTGAAGTTTTCTGGGTC | 52 | 53.8 | GG | 0.92 | 0.87 | 330 | 330 | 330 |
| GAAGGATATTTGGCCTCA | GCCCACACATGCCGAAAC | 52 | 60.8 | GG | 0.98 | 0.69 | 186 | 186 | 186 |
| GCCCACACATGCCGAAAC | TACGAAGGCAGTTGCTAT | 60.8 | 53.9 | GG | 0.98 | 0.67 | 153 | 153 | 153 |
| CTACGAAGGCAGTTGCTA | GCCCACACATGCCGAAAC | 54.9 | 60.8 | GG | 0.98 | 0.67 | 154 | 154 | 154 |
| CCTACGAAGGCAGTTGCT | GCCCACACATGCCGAAAC | 57.8 | 60.8 | GG | 0.98 | 0.67 | 155 | 155 | 155 |
| CCCACACATGCCGAAACG | TACGAAGGCAGTTGCTAT | 60.5 | 53.9 | GG | 0.98 | 0.67 | 152 | 152 | 152 |
| CCCACACATGCCGAAACG | CCTACGAAGGCAGTTGCT | 60.5 | 57.8 | GG | 0.98 | 0.67 | 154 | 154 | 154 |
| CACATGCCGAAACGTACA | TACGAAGGCAGTTGCTAT | 56.6 | 53.9 | GG | 0.90 | 0.64 | 148 | 148 | 148 |
| CACATGCCGAAACGTACA | CCTACGAAGGCAGTTGCT | 56.6 | 57.8 | GG | 0.90 | 0.64 | 150 | 150 | 150 |
| **Marker** | **12S** | | | | | | | | |
| AAAGACTTAGTCCTAACC | ACAAGATTTACCAACCCT | 48.4 | 51.1 | GG | 1.00 | 0.92 | 207 | 213 | 210.68 |
| AAAGACTTAGTCCTAACC | CACAAGATTTACCAACCC | 48.4 | 51.4 | GG | 1.00 | 0.92 | 208 | 214 | 211.68 |
| AAAGACTTAGTCCTAACC | GCTGGCACAAGATTTACC | 48.4 | 54.7 | GG | 1.00 | 0.92 | 213 | 219 | 216.68 |
| AAAGACTTAGTCCTAACC | GGCTGGCACAAGATTTAC | 48.4 | 54.7 | GG | 1.00 | 0.92 | 214 | 220 | 217.68 |
| AAGACTTAGTCCTAACCT | ACAAGATTTACCAACCCT | 49.1 | 51.1 | GG | 1.00 | 0.92 | 206 | 212 | 209.68 |
| AAGACTTAGTCCTAACCT | CACAAGATTTACCAACCC | 49.1 | 51.4 | GG | 1.00 | 0.92 | 207 | 213 | 210.68 |
| AAGACTTAGTCCTAACCT | GCTGGCACAAGATTTACC | 49.1 | 54.7 | GG | 1.00 | 0.92 | 212 | 218 | 215.68 |
| AAGACTTAGTCCTAACCT | GGCTGGCACAAGATTTAC | 49.1 | 54.7 | GG | 1.00 | 0.92 | 213 | 219 | 216.68 |
| ACAAGATTTACCAACCCT | AGACTTAGTCCTAACCTT | 51.1 | 49.1 | GG | 1.00 | 0.92 | 205 | 211 | 208.68 |
| AGACTTAGTCCTAACCTT | CACAAGATTTACCAACCC | 49.1 | 51.4 | GG | 1.00 | 0.92 | 206 | 212 | 209.68 |
| AGACTTAGTCCTAACCTT | GCTGGCACAAGATTTACC | 49.1 | 54.7 | GG | 1.00 | 0.92 | 211 | 217 | 214.68 |
| AGACTTAGTCCTAACCTT | GGCTGGCACAAGATTTAC | 49.1 | 54.7 | GG | 1.00 | 0.92 | 212 | 218 | 215.68 |
| AAACTGGGATTAGATACC | TACTGCTAAATCCGCCTT | 48.5 | 53.9 | GG | 1.00 | 0.92 | 337 | 344 | 340.44 |

| | | | | | | | | | |
|---|---|---|---|---|---|---|---|---|---|
| AAACTGGGATTAGATACC | TTACTGCTAAATCCGCCT | 48.5 | 53.9 | GG | 1.00 | 0.92 | 338 | 345 | 341.44 |
| AACTGGGATTAGATACCC | TACTGCTAAATCCGCCTT | 50.5 | 53.9 | GG | 1.00 | 0.92 | 336 | 343 | 339.44 |
| AACTGGGATTAGATACCC | TTACTGCTAAATCCGCCT | 50.5 | 53.9 | GG | 1.00 | 0.92 | 337 | 344 | 340.44 |
| ACTGGGATTAGATACCCC | TTACTGCTAAATCCGCCT | 52.6 | 53.9 | GG | 1.00 | 0.92 | 336 | 343 | 339.44 |
| AAAGACTTAGTCCTAACC | CTGGCACAAGATTTACCA | 48.4 | 53 | GG | 0.98 | 0.92 | 212 | 218 | 215.67 |
| AAGACTTAGTCCTAACCT | CTGGCACAAGATTTACCA | 49.1 | 53 | GG | 0.98 | 0.92 | 211 | 217 | 214.67 |
| AGACTTAGTCCTAACCTT | CTGGCACAAGATTTACCA | 49.1 | 53 | GG | 0.98 | 0.92 | 210 | 216 | 213.67 |
| AAACTGGGATTAGATACC | ACTGCTAAATCCGCCTTC | 48.5 | 55.5 | GG | 0.98 | 0.92 | 336 | 343 | 339.41 |
| AACTGGGATTAGATACCC | ACTGCTAAATCCGCCTTC | 50.5 | 55.5 | GG | 0.98 | 0.92 | 335 | 342 | 338.41 |
| AAAGACTTAGTCCTAACC | AATGTTAATTACTGCTGA | 48.4 | 46.7 | GG | 1.00 | 0.90 | 148 | 155 | 152.74 |
| AAAGACTTAGTCCTAACC | TAATGTTAATTACTGCTG | 48.4 | 45 | GG | 1.00 | 0.90 | 149 | 156 | 153.74 |
| AAAGACTTAGTCCTAACC | TTAATGTTAATTACTGCT | 48.4 | 43.9 | GG | 1.00 | 0.90 | 150 | 157 | 154.74 |
| AAAGACTTAGTCCTAACC | CTTAATGTTAATTACTGC | 48.4 | 43.9 | GG | 1.00 | 0.90 | 151 | 158 | 155.74 |
| AAAGACTTAGTCCTAACC | GCTTAATGTTAATTACTG | 48.4 | 43.9 | GG | 1.00 | 0.90 | 152 | 159 | 156.74 |
| AAAGACTTAGTCCTAACC | TGCTTAATGTTAATTACT | 48.4 | 43.9 | GG | 1.00 | 0.90 | 153 | 160 | 157.74 |
| AAAGACTTAGTCCTAACC | TTGCTTAATGTTAATTAC | 48.4 | 43.2 | GG | 1.00 | 0.90 | 154 | 161 | 158.74 |
| AAAGACTTAGTCCTAACC | ATTGCTTAATGTTAATTA | 48.4 | 41.8 | GG | 1.00 | 0.90 | 155 | 162 | 159.74 |
| AAGACTTAGTCCTAACCT | AATGTTAATTACTGCTGA | 49.1 | 46.7 | GG | 1.00 | 0.90 | 147 | 154 | 151.74 |
| AAGACTTAGTCCTAACCT | TAATGTTAATTACTGCTG | 49.1 | 45 | GG | 1.00 | 0.90 | 148 | 155 | 152.74 |
| AAGACTTAGTCCTAACCT | TTAATGTTAATTACTGCT | 49.1 | 43.9 | GG | 1.00 | 0.90 | 149 | 156 | 153.74 |
| AAGACTTAGTCCTAACCT | CTTAATGTTAATTACTGC | 49.1 | 43.9 | GG | 1.00 | 0.90 | 150 | 157 | 154.74 |
| AAGACTTAGTCCTAACCT | GCTTAATGTTAATTACTG | 49.1 | 43.9 | GG | 1.00 | 0.90 | 151 | 158 | 155.74 |
| AAGACTTAGTCCTAACCT | TGCTTAATGTTAATTACT | 49.1 | 43.9 | GG | 1.00 | 0.90 | 152 | 159 | 156.74 |
| AAGACTTAGTCCTAACCT | TTGCTTAATGTTAATTAC | 49.1 | 43.2 | GG | 1.00 | 0.90 | 153 | 160 | 157.74 |
| AAGACTTAGTCCTAACCT | ATTGCTTAATGTTAATTA | 49.1 | 41.8 | GG | 1.00 | 0.90 | 154 | 161 | 158.74 |
| AATGTTAATTACTGCTGA | AGACTTAGTCCTAACCTT | 46.7 | 49.1 | GG | 1.00 | 0.90 | 146 | 153 | 150.74 |
| AGACTTAGTCCTAACCTT | TAATGTTAATTACTGCTG | 49.1 | 45 | GG | 1.00 | 0.90 | 147 | 154 | 151.74 |
| AGACTTAGTCCTAACCTT | TTAATGTTAATTACTGCT | 49.1 | 43.9 | GG | 1.00 | 0.90 | 148 | 155 | 152.74 |
| AGACTTAGTCCTAACCTT | CTTAATGTTAATTACTGC | 49.1 | 43.9 | GG | 1.00 | 0.90 | 149 | 156 | 153.74 |
| AGACTTAGTCCTAACCTT | GCTTAATGTTAATTACTG | 49.1 | 43.9 | GG | 1.00 | 0.90 | 150 | 157 | 154.74 |
| AGACTTAGTCCTAACCTT | TGCTTAATGTTAATTACT | 49.1 | 43.9 | GG | 1.00 | 0.90 | 151 | 158 | 155.74 |
| AGACTTAGTCCTAACCTT | TTGCTTAATGTTAATTAC | 49.1 | 43.2 | GG | 1.00 | 0.90 | 152 | 159 | 156.74 |
| AGACTTAGTCCTAACCTT | ATTGCTTAATGTTAATTA | 49.1 | 41.8 | GG | 1.00 | 0.90 | 153 | 160 | 157.74 |
| **Marker** | **Dloop** | | | | | | | | |
| ACCCCACCTTCGGCATGG | ACTATTATTCATATATAT | 62.8 | 35.7 | GG | 1.00 | 1.00 | 101 | 101 | 101 |
| ACTATTATTCATATATAT | GACCCCACCTTCGGCATG | 35.7 | 61.5 | GG | 1.00 | 1.00 | 102 | 102 | 102 |
| ACTATTATTCATATATAT | AGACCCCACCTTCGGCAT | 35.7 | 61.2 | GG | 1.00 | 1.00 | 103 | 103 | 103 |
| ACTATTATTCATATATAT | GAGACCCCACCTTCGGCA | 35.7 | 62.1 | GG | 1.00 | 1.00 | 104 | 104 | 104 |
| ACTATTATTCATATATAT | AGAGACCCCACCTTCGGC | 35.7 | 61.8 | GG | 1.00 | 1.00 | 105 | 105 | 105 |
| ACTATTATTCATATATAT | CAGAGACCCCACCTTCGG | 35.7 | 60.1 | GG | 1.00 | 1.00 | 106 | 106 | 106 |
| ACAGAGACCCCACCTTCG | ACTATTATTCATATATAT | 59 | 35.7 | GG | 1.00 | 1.00 | 107 | 107 | 107 |
| ACTATTATTCATATATAT | AGGGAAATTCTATTGATA | 35.7 | 44.7 | GG | 1.00 | 1.00 | 126 | 126 | 126 |
| ACTATTATTCATATATAT | TAGGGAAATTCTATTGAT | 35.7 | 44.7 | GG | 1.00 | 1.00 | 127 | 127 | 127 |
| ACTATTATTCATATATAT | GTAGGGAAATTCTATTGA | 35.7 | 46.1 | GG | 1.00 | 1.00 | 128 | 128 | 128 |
| ACTATTATTCATATATAT | CGTAGGGAAATTCTATTG | 35.7 | 48 | GG | 1.00 | 1.00 | 129 | 129 | 129 |
| ACTATTATTCATATATAT | CCGTAGGGAAATTCTATT | 35.7 | 48.9 | GG | 1.00 | 1.00 | 130 | 130 | 130 |
| ACTATTATTCATATATAT | TCCGTAGGGAAATTCTAT | 35.7 | 49.7 | GG | 1.00 | 1.00 | 131 | 131 | 131 |
| ACTATTATTCATATATAT | ATCCGTAGGGAAATTCTA | 35.7 | 49.7 | GG | 1.00 | 1.00 | 132 | 132 | 132 |

| | | | | | | | | | |
|---|---|---|---|---|---|---|---|---|---|
| ACTATTATTCATATATAT | TATCCGTAGGGAAATTCT | 35.7 | 49.7 | GG | 1.00 | 1.00 | 133 | 133 | 133 |
| ACTATTATTCATATATAT | ATATCCGTAGGGAAATTC | 35.7 | 48.7 | GG | 1.00 | 1.00 | 134 | 134 | 134 |
| AATATCCGTAGGGAAATT | ACTATTATTCATATATAT | 48 | 35.7 | GG | 1.00 | 1.00 | 135 | 135 | 135 |
| ACTATTATTCATATATAT | GAATATCCGTAGGGAAAT | 35.7 | 48.7 | GG | 1.00 | 1.00 | 136 | 136 | 136 |
| ACTATTATTCATATATAT | AGAATATCCGTAGGGAAA | 35.7 | 49.7 | GG | 1.00 | 1.00 | 137 | 137 | 137 |
| AAGAATATCCGTAGGGAA | ACTATTATTCATATATAT | 49.7 | 35.7 | GG | 1.00 | 1.00 | 138 | 138 | 138 |
| ACTATTATTCATATATAT | CAAGAATATCCGTAGGGA | 35.7 | 50.8 | GG | 1.00 | 1.00 | 139 | 139 | 139 |
| ACTATTATTCATATATAT | CCAAGAATATCCGTAGGG | 35.7 | 52.1 | GG | 1.00 | 1.00 | 140 | 140 | 140 |
| ACTATTATTCATATATAT | CCCAAGAATATCCGTAGG | 35.7 | 52.1 | GG | 1.00 | 1.00 | 141 | 141 | 141 |
| ACTATTATTCATATATAT | CCCCAAGAATATCCGTAG | 35.7 | 52.1 | GG | 1.00 | 1.00 | 142 | 142 | 142 |
| ACTATTATTCATATATAT | TCCCCAAGAATATCCGTA | 35.7 | 52.1 | GG | 1.00 | 1.00 | 143 | 143 | 143 |
| ACTATTATTCATATATAT | GTCCCCAAGAATATCCGT | 35.7 | 54.1 | GG | 1.00 | 1.00 | 144 | 144 | 144 |
| ACTATTATTCATATATAT | TGTCCCCAAGAATATCCG | 35.7 | 54.1 | GG | 1.00 | 1.00 | 145 | 145 | 145 |
| ACTATTATTCATATATAT | TTGTCCCCAAGAATATCC | 35.7 | 51.5 | GG | 1.00 | 1.00 | 146 | 146 | 146 |
| ACTATTATTCATATATAT | TTTGTCCCCAAGAATATC | 35.7 | 49.5 | GG | 1.00 | 1.00 | 147 | 147 | 147 |
| ACTATTATTCATATATAT | GTTTGTCCCCAAGAATAT | 35.7 | 49.8 | GG | 1.00 | 1.00 | 148 | 148 | 148 |
| ACTATTATTCATATATAT | AGTTTGTCCCCAAGAATA | 35.7 | 50.8 | GG | 1.00 | 1.00 | 149 | 149 | 149 |
| ACTATTATTCATATATAT | GAGTTTGTCCCCAAGAAT | 35.7 | 52.5 | GG | 1.00 | 1.00 | 150 | 150 | 150 |
| ACTATTATTCATATATAT | AGAGTTTGTCCCCAAGAA | 35.7 | 53.4 | GG | 1.00 | 1.00 | 151 | 151 | 151 |
| ACTATTATTCATATATAT | CAGAGTTTGTCCCCAAGA | 35.7 | 54.5 | GG | 1.00 | 1.00 | 152 | 152 | 152 |
| ACAGAGTTTGTCCCCAAG | ACTATTATTCATATATAT | 54.8 | 35.7 | GG | 1.00 | 1.00 | 153 | 153 | 153 |
| ACTATTATTCATATATAT | TACAGAGTTTGTCCCCAA | 35.7 | 53.1 | GG | 1.00 | 1.00 | 154 | 154 | 154 |
| ACTATTATTCATATATAT | ATACAGAGTTTGTCCCCA | 35.7 | 52.9 | GG | 1.00 | 1.00 | 155 | 155 | 155 |
| ACTATTATTCATATATAT | CATACAGAGTTTGTCCCC | 35.7 | 52.9 | GG | 1.00 | 1.00 | 156 | 156 | 156 |
| ACTATTATTCATATATAT | CCATACAGAGTTTGTCCC | 35.7 | 52.9 | GG | 1.00 | 1.00 | 157 | 157 | 157 |
| ACCATACAGAGTTTGTCC | ACTATTATTCATATATAT | 51.9 | 35.7 | GG | 1.00 | 1.00 | 158 | 158 | 158 |
| ACTATTATTCATATATAT | TACCATACAGAGTTTGTC | 35.7 | 48.9 | GG | 1.00 | 1.00 | 159 | 159 | 159 |
| ACTATTATTCATATATAT | CTACCATACAGAGTTTGT | 35.7 | 48.9 | GG | 1.00 | 1.00 | 160 | 160 | 160 |
| ACTATTATTCATATATAT | GCTACCATACAGAGTTTG | 35.7 | 50.7 | GG | 1.00 | 1.00 | 161 | 161 | 161 |
| ACTATTATTCATATATAT | GGCTACCATACAGAGTTT | 35.7 | 51.7 | GG | 1.00 | 1.00 | 162 | 162 | 162 |
| ACTATTATTCATATATAT | CGGCTACCATACAGAGTT | 35.7 | 54.2 | GG | 1.00 | 1.00 | 163 | 163 | 163 |
| ACTATTATTCATATATAT | CCGGCTACCATACAGAGT | 35.7 | 56.2 | GG | 1.00 | 1.00 | 164 | 164 | 164 |
| ACTATTATTCATATATAT | AGATAACCTGGTCCGACA | 35.7 | 55.1 | GG | 1.00 | 1.00 | 218 | 218 | 218 |
| ACTATTATTCATATATAT | GAGATAACCTGGTCCGAC | 35.7 | 54.8 | GG | 1.00 | 1.00 | 219 | 219 | 219 |
| ACTATTATTCATATATAT | AGAGATAACCTGGTCCGA | 35.7 | 54.5 | GG | 1.00 | 1.00 | 220 | 220 | 220 |
| AAGAGATAACCTGGTCCG | ACTATTATTCATATATAT | 53.7 | 35.7 | GG | 1.00 | 1.00 | 221 | 221 | 221 |
| ACTATTATTCATATATAT | TAAGAGATAACCTGGTCC | 35.7 | 50.2 | GG | 1.00 | 1.00 | 222 | 222 | 222 |

Tm: melting temperature of primers; Quality: G (good) / B (Bad); Bc: coverage index (probability of amplification success); Bs: specificity index (taxon discrimination power).

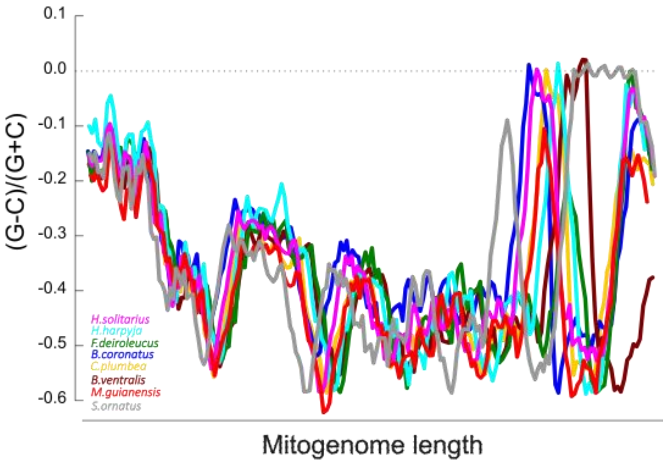


Figure S1. Nucleotide skew across the complete mitogenome of eight Neotropical raptor species.
